# Single cell transcriptome sequencing on the Nanopore platform with ScNapBar

**DOI:** 10.1101/2020.10.16.342626

**Authors:** Qi Wang, Sven Bönigk, Volker Böhm, Niels Gehring, Janine Altmüller, Christoph Dieterich

## Abstract

The current ecosystem of single cell RNA-seq platforms is rapidly expanding, but robust solutions for single cell and single molecule full-length RNA sequencing are virtually absent. A high-throughput solution that covers all aspects is necessary to study the complex life of mRNA on the single cell level. The Nanopore platform offers long read sequencing and can be integrated with the popular single cell sequencing method on the 10x Chromium platform. However, the high error-rate of Nanopore reads poses a challenge in downstream processing (e.g. for cell barcode assignment). We propose a solution to this particular problem by using a hybrid sequencing approach on Nanopore and Illumina platforms. Our software ScNapBar enables cell barcode assignment with high accuracy, especially if sequencing saturation is low. ScNapBar uses unique molecular identifier (UMI) or Naïve Bayes probabilistic approaches in the barcode assignment, depending on the available Illumina sequencing depth. We have benchmarked the two approaches on simulated and real Nanopore datasets. We further applied ScNapBar to pools of cells with an active or a silenced non-sense mediated RNA decay pathway. Our Nanopore read assignment distinguishes the respective cell populations and reveals characteristic nonsense-mediated mRNA decay events depending on cell status.

## INTRODUCTION

Full-length cDNA sequencing allows us to investigate the differential iso-forms of transcripts, which is especially useful in studying the complex life of mRNA. Compared to the Illumina sequencing approaches, third-generation sequencing generates much longer reads and thus avoids artifacts from transcriptome assembly, but often has limitations such as low throughput and poor base-calling accuracy. Two principal third-generation sequencing platforms exist: Oxford Nanopore Technologies (ONT) and Pacific Biosciences (PacBio) (Volden et al., 2018). Others and we chose the ONT platform to study full-length mRNA transcripts due to its better scalability and flexibility (Lebrigand et al., 2020). Full-length transcriptome sequencing can be taken to the single level by sequencing barcoded 10x Genomics cDNA libraries. However, this brings about certain challenges, which we address in our work.

First, the native error rate of Nanopore DNA sequencing is < 5% on the latest R10.3 platform (http://nanoporetech.com) as opposed to the typical Illumina error rate of 0.1%. Due to its high error rate, barcode identification and assignment are challenging for single-cell sequencing. In the 10X Genomics single-cell protocol, about 99% barcode sequences from Illumina sequencing can be exactly matched to the 16-bp cell barcodes, while with Nanopore sequencing, the exact matches are less than 50% (0.999^16^ vs. 0.95^16^). Many experimental and computational approaches have been developed to correct Nanopore data. For example, the rolling circle to concate-meric consensus (R2C2) approach can produce two million full-length cDNA sequences per MinION flow cell and achieved 98% accuracy (Volden et al., 2018; Cole et al., 2020; Volden and Vollmers, 2020). Single-cell Nanopore sequencing with UMIs (ScNaUmi-seq) can assign cellular barcode with 99.8% accuracy (Lebrigand et al., 2020). However, R2C2 requires sufficient sequencing coverage to call consensus reads, and ScNaUmi-seq requires high sequencing depth to guarantee an adequate overlap of UMI sequences between Illumina and Nanopore libraries.

On the other hand, end-to-end solutions for barcode demultiplexing and read quality filtering on the ONT platform are still in its infancy. For example, Mandalorion uses BLAT (Kent, 2002) for barcode demultiplexing (Byrne et al., 2017). Porechop (https://github.com/rrwick/Porechop) uses SeqAn (Döring et al., 2008) for adapter removal and barcode demultiplexing in Nanopore sequencing, but it is based on the best alignment which could be error-prone. Minibar (Krehenwinkel et al., 2019), Deep-binner (Wick et al., 2018), and DeePlexiCon (Smith et al., 2020) are only suitable for multiplexing a few barcoded samples rather than the single-cell library which contains several thousands of barcodes.

Therefore, we developed a software tool called ScNapBar (single-cell Nanopore barcode demultiplexer) that demultiplexes Nanopore barcodes and is particularly suited for low depth Illumina and Nanopore sequencing. We evaluated the performance of ScNapBar and demonstrated its high accuracy in cell barcode assignment for simulated and real Nanopore data. Our workflow is presented in Fig. 1.

**Figure 1:**
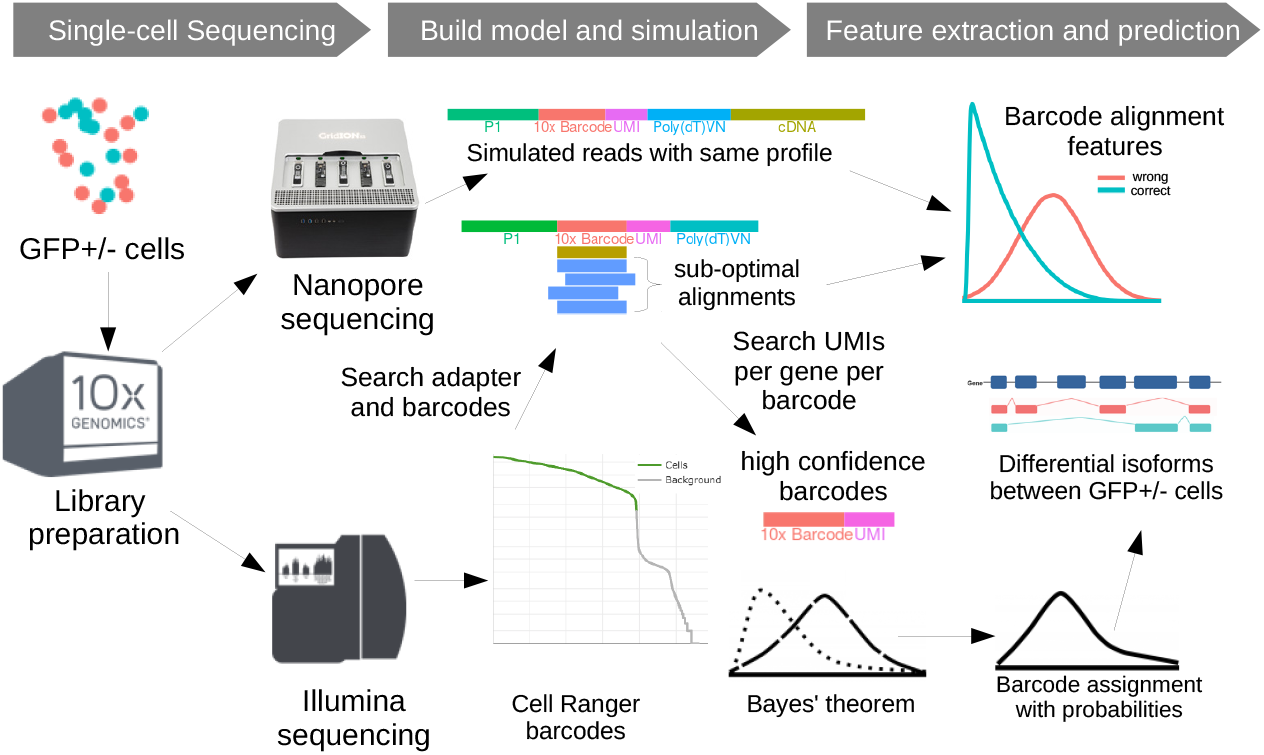
Combined Single-cell Illumina and Nanopore sequencing strategy. GFP+/- cells are pooled and sequenced on the Illumina and Nanopore platform. The Nanopore platform generates long cDNA sequencing read that are used in barcode calling and estimating read error parameters. The Illumina data are used to estimate the total number of cells in sequencing and the represented cell barcodes. The simulated data are then used to parameterize a Bayesian model of barcode alignment features to discriminate correct vs. false barcode assignments. This model is then used on the real data to assign cell barcodes to Nanopore reads. The GFP label and known NMD transcripts can be used to validate this assignment.

## RESULTS

### Benchmarking the two ScNapBar run modes

ScNapBar offers to run modes. The first one uses cell barcode and UMI information without any additional modeling aspect. The second one introduces a probabilistic model, which performs very well in cases of low sequencing saturation (i.e. UMI coverage in Illumina data).

#### The UMI approach of ScNapBar

The UMI approach requires a matching cell barcodes and UMI tag and was first developed in Sicelore (Lebrigand et al., 2020). Any cell barcode predictions that are supported by the presence of both, barcode and UMI alignment, are very reliable. We performed an *in silico* benchmark of cell barcode assignment when both, cell barcode and UMI, are found in the Nanopore read. We observed an average specificity of 99.9% (ScNapBar) and 99.8% (Sicelore) over 100 averaged simulation runs (Fig. 2a). As expected, sensitivity heavily depends on Illumina sequencing saturation (Fig. 2a). As the UMI approach relies on consistent genomic mappings for the Illumina and Nanopore reads, other challenges include: insufficient or inaccurate genome annotations causing wrong gene assignment; chimeric or super-long Nanopore reads assigned to multiple genes increase the risk of assigning a false UMI.

**Figure 2:**
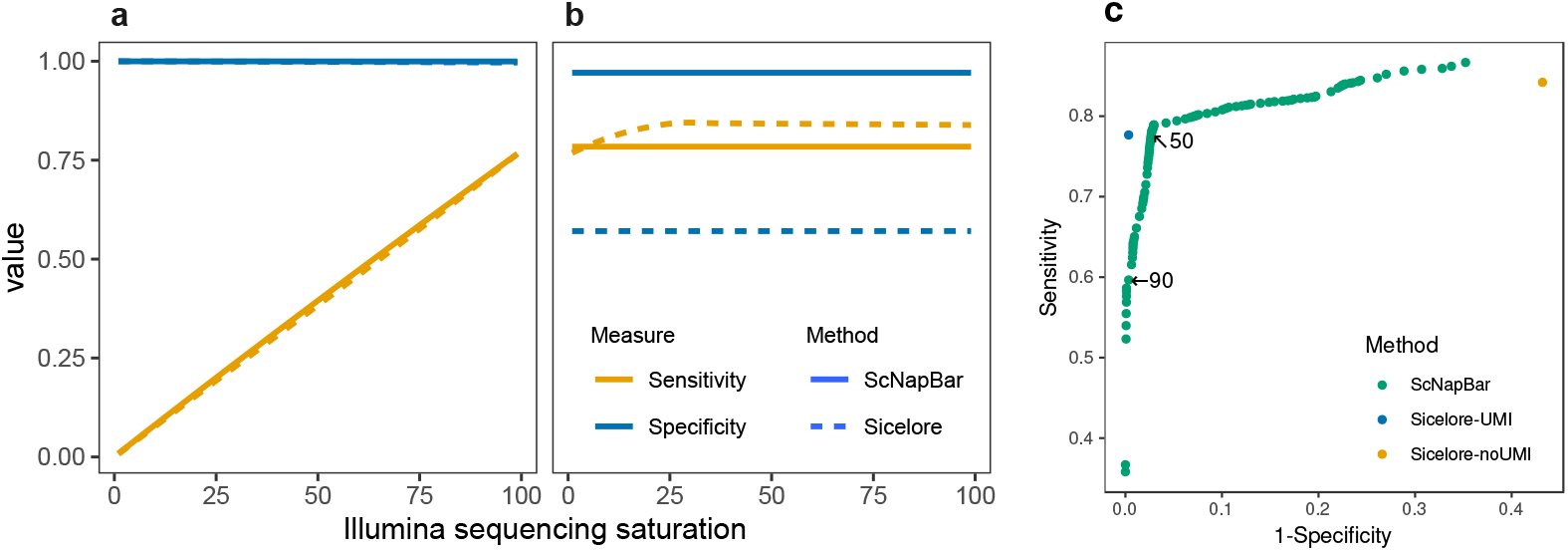
Sensitivity and specificity of ScNapBar and Sicelore on 100 Illumina libraries with different levels of saturation. (a) Barcode assignment with UMI matches. (b) Barcode assignment without UMI matches (ScNapBar score >50). (c) Benchmark of the specificity and sensitivity of the Illumina library with 100% saturation. We compared the barcode assignments with ScNapBar score >1-99, and the assignments from Sicelore with UMI support are roughly equivalent to the ScNapBar score >90.

#### The probabilistic approach of ScNapBar

Complementary to the UMI approach, we implemented a Bayesian approach in ScNapBar, which covers the situation of low Illumina sequencing saturation. In our second approach, UMI alignments are no longer used. ScNapBar evaluates probability scores for each barcode alignment instead. Illumina sequencing saturation measures the uniqueness of the transcripts detected in the Illumina library. Given that we have performed Illumina and Nanopore sequencing in our approach, the Illumina sequencing saturation limits the overlap of cell barcodes and UMIs with the low depth Nanopore libraries. To explore more realistic saturation scenarios, we estimated the Illumina sequencing saturation for our pilot data set with the Cell Ranger software. Herein, sequencing saturation is calculated as

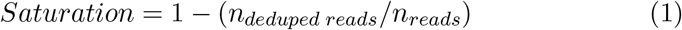

where *n_deduped reads_* is the number of unique (valid cell-barcode, valid UMI, gene) combinations among confidently mapped reads and *n_reads_* is the total number of confidently mapped, valid cell-barcode, valid UMI reads. For example, we have observed a saturation of 11.3% for our pilot data set.

We have simulated one million Nanopore reads with an error model, which was estimated from our reference Nanopore libraries (see Methods) using the same gene-barcode-UMI composition as given by the Illumina library and a sequencing saturation of 100%. We trained a Naïve Bayes classifier (see Methods) from barcode and adapter alignments of one Nanopore library, and applied the model for computing the likelihood of the matched barcodes *P*(*r|b_i_*) on the other library. Then we used the frequencies of the given barcodes in the Illumina library as prior probabilities *P*(*b_i_*), and calculated the posterior probability *P*(*b_i_|r*) from the likelihood and prior probabilities. We scored each barcode alignment by multiplying the *P*(*b_i_|r*) by 100, and assigned the best matching barcode with the highest score (> 50) as predicted barcode assignment. Using the probability scores as mentioned, ScNapBar correctly assigned 65.8% barcodes from one million simulated Nanopore reads, of which 26.5% contains at least one mismatch or indel (Suppl. Fig. S1).

We estimate a user data specific error model, simulate data from which users pick the Bayes score cutoff, which meets their requirements on sensitivity and specificity, respectively. We inspected the densities of the probability scores by examining the ground-truth barcodes, and confirmed that the correct barcode assignments are enriched in high scoring barcodes (Suppl. Fig. S2b).

Our probabilistic model outperforms Sicelore for cases where UMI information is sparse and cannot be used to assign cell barcodes. In the absence of UMIs, ScNapBar reaches 97.1% specificity while Sicelore only reaches only 57.1% (Fig. 2b).

We examined performance metrics of cell barcode assignment over a range of score cutoffs (from 1 to 99), and the specificity increases while the sensitivity decreases along with the increased thresholds (Suppl. Fig. S3). We pooled the simulated results from FC1 and FC2 together, and use the Sicelore assignments as baselines. As some cutoff thresholds, ScNapBar has better F1 scores than Sicelore (e.g., cutoff=50), and ScNapBar score >90 is as accurate as Sicelore with UMI from the Receiver-Operating Characteristic (ROC) graph (Fig. 2c).

#### The runtime performance of ScNapBar

ScNapBar is based on the Needleman-Wunsch algorithm (gap-end free, semi-global sequence alignment) of FLEXBAR (Dodt et al., 2012; Roehr et al., 2017) and Sicelore is based on the “brute force approach” which hashes all possible sequence tag variants (including indels) up to a certain edit distance (2 or 3) of the given barcode sequences. The time complexity of ScNapBar and Sicelore can be represented as Eq. 2a and Eq. 2b, respectively.

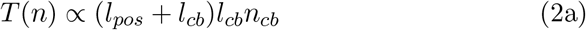

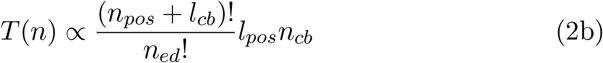

where *n_pos_* is the number of nucleotides downstream of the adapter, and *l_pos_* = *2n_pos_* +1 as Sicelore typically searches the same number of nucleotides upstream and downstream of the ending position of the adapter. *n_cb_* stands for the number of barcodes in the whitelist from Illumina sequencing. *n_ed_* is typically two or three as larger edit distances increase runtime drastically and are not necessary due to the increasing error rate. *l_cb_* is the length of the barcode and is 16 in this study.

We compared the runtime between ScNapBar and Sicelore with regards to start positions of barcodes (number of nucleotides between adapter and barcode). We discovered that Sicelore may be orders of magnitude slower than ScNapBar given the same search space (2,052 cellular barcodes, edit distance=3), but also its runtime increases exponentially as the barcode start position increases(Fig. 3b). Therefore, the default setting in Sicelore only searches ± 1-nt from the end of the adapter, which may limit the nucleotides to search and cause false positives. We created 2×2 contingency tables of the number of correct and false assignments caused by various factors (e.g., indels ≽ 3 against < 3), and performed Fisher’s test. The results showed that the odds ratio of “barcode start position ≽ 3” from Sicelore is 24.8, while the odds ratio of the same test from ScNapBar is only 0.14 (Suppl. Table S1). This implies allowing more nucleotides from the start of the barcode can effectively reduce the false-positive rate, which is feasible using less time with ScNapBar.

**Figure 3:**
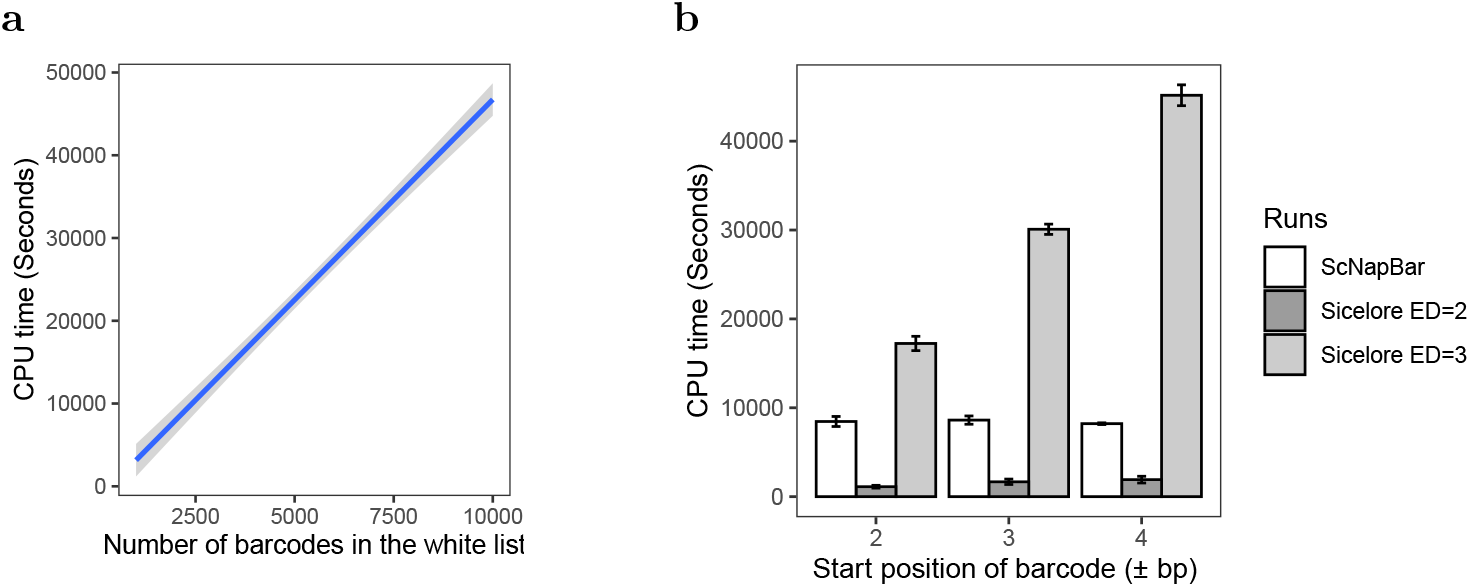
Sicelore and ScNapBar CPU time comparison. (a) ScNapBar CPU time depends on the number of whitelist barcodes (allowing an edit distance of >2 and and offset of up to 4bp between adapter and barcode). Gray area represents the standard deviation for 10 runs. (b) Comparison of ScNapBar and Sicelore CPU times. Benchmark was measured using one million barcode sequences and 2,052 barcodes in the whitelist.

We also performed real runtime comparison on barcode assignment on the previously simulated one million Nanopore reads. In this test, we provided ScNapBar ten barcode white lists which contain from 1,000 to 10,000 most abundant barcodes, and ScNapBar’s runtime is only dependent on the number of barcodes to search given the other factors are fixed in this study (Fig. 3a). Then we tested Sicelore with searching parameters of barcode edit distance between two and three, barcode start position from ±2 bp to ±4 bp, and UMI edit distance of 0. ScNapBar requires only one-fifth CPU time than Sicelore when ±4 bp barcode start position and three barcode edit distance are considered in both programs (Fig. 3b).

### The performance of ScNapBar on the real data

#### The performance of ScNapBar on an Illumina library with high sequencing saturations

We tested our ScNapBar software with the UMI approach (option 1) on the dataset from the Sicelore paper (NCBI GEO GSE130708). Herein, Illumina sequencing saturation reaches 90.5%. We extracted the UMI whitelists for each gene or genomic window (500bp) from the Illumina library, and set the minimum length of UMI match to 7 in ScNapBar. Sicelore and ScNapBar assigned barcodes to 84.3% and 77.2% of the 9,743,819 Nanopore reads (Suppl. Fig. S4), respectively. 88.4% of the assigned barcodes are identical.

#### The performance of ScNapBar on an Illumina library with low sequencing saturations

We ran ScNapBar with the Bayesian approach (option 2) on our NMD dataset, which only has an Illumina saturation of 11.3%. ScNapBar assigns 35.0% and 36.3% of the Nanopore reads to cell barcodes with probability score >50, while Sicelore assigns 40.8% and 42.5% without using UMIs (“Assigned to barcode” in Fig. 5) and only assigns 4.0% and 4.2% of the Nanopore reads using the UMI approach for FC1 and FC2, respectively. Based on our previous simulations, we estimate that a greater proportion (also by absolute numbers) of ScNapBar assignments are correct (“Correctly assigned” in Fig. 5).

**Figure 4:**
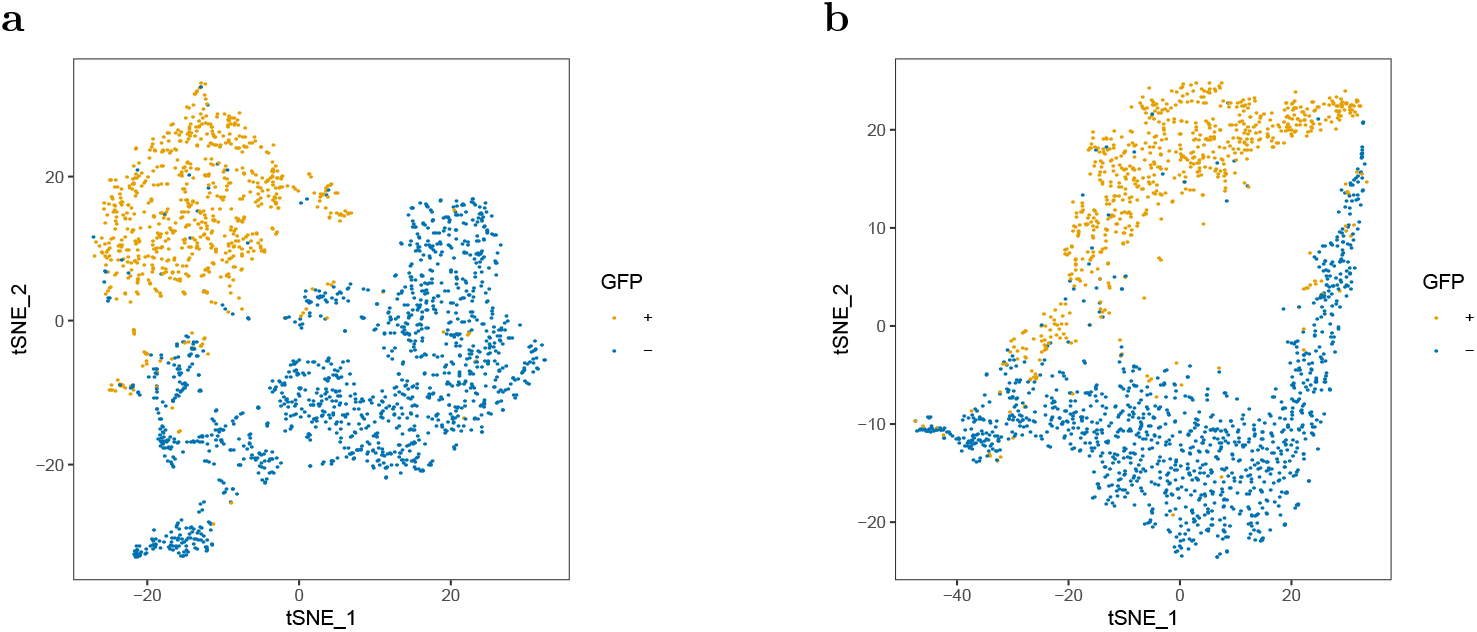
The t-SNE plots of gene-cell matrices. (a) Illumina. (b) Nanopore.

**Figure 5:**
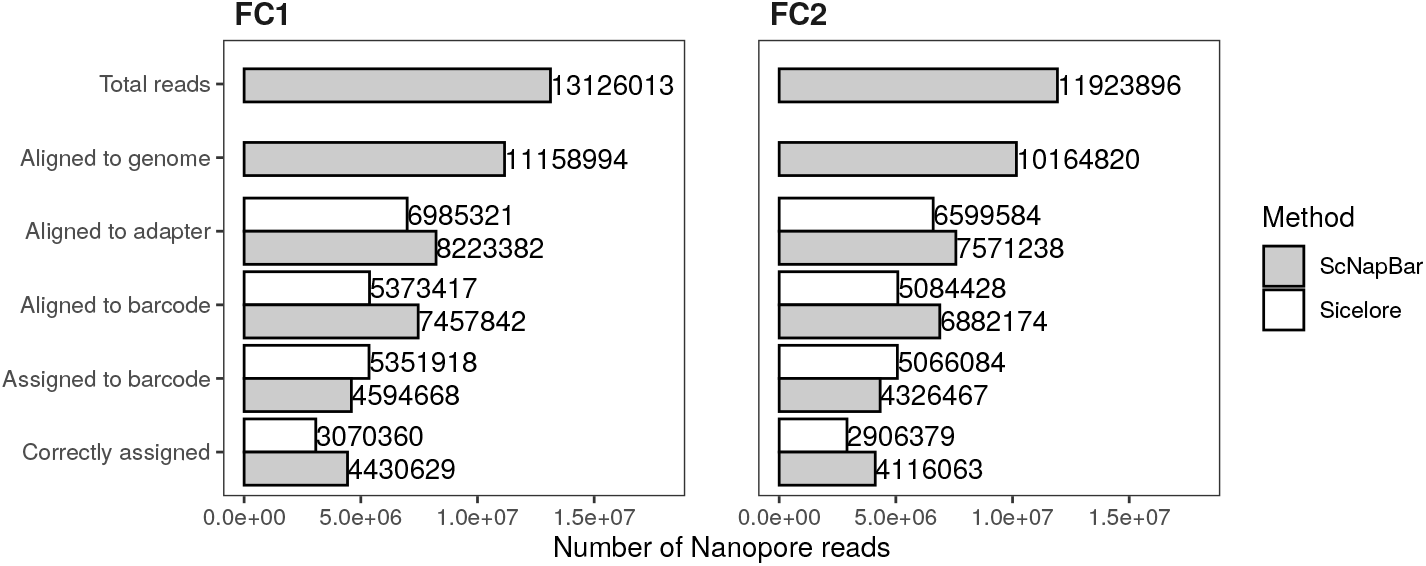
Number of the Nanopore reads identified by ScNapBar and Sicelore from each step. The number of the correctly assigned reads is calculated from the specificity of the assignment in the simulation.

#### Single cell clustering and splicing in a pool of wildtype and NMD mutant cells

Although alternative splicing increases the coding potential of the human genome, aberrant isoforms are frequently generated that contain premature termination codons (PTCs) (Lewis et al., 2003). Regular stop codons are normally located in the last exon of a transcript or at least 50 nucleotides upstream of the last exon-exon junction (Lindeboom et al., 2019). Alternative splicing can result in PTCs by exon inclusion/exclusion events or can convert normal stop codons into PTCs by splicing in the 3’ UTR. Transcripts harboring PTCs are rapidly degraded by the nonsense-mediated mRNA decay (NMD) machinery, not only to remove faulty mRNAs, but also to fine-tune and regulate the transcriptome. 5-40% of all expressed human genes are directly or indirectly altered in expression levels, splicing pattern, or isoform composition by the NMD pathway (Boehm et al., 2020). We have sequenced a pool of NMD active and inactive cells and expect to see an enrichment of transcripts with PTCs in GFP-cells.

We use the GFP label as an independent confirmation of cellular NMD status and pooled data from both experiments (FC1 and FC2). For the Nanopore data, Seurat identifies 13,807 expressed genes across 1,850 cells. We extracted the GFP+ barcodes from the Illumina reads mapping, and rendered the corresponding cells in different colors in the t-SNE plots (Fig. 4). The locations of the GFP+ cells appear in distinct sub-clusters in the Illumina and Nanopore t-SNE plots.

We characterized the structural changes of the assembled Nanopore transcripts based on our customized transcriptome annotations using NMD Classifier (Hsu et al., 2017). The pool of SMG7-KO/SMG6-KD (GFP-) cells harbors almost twice as many inclusion/exclusion events, which lead to the formation of a PTC (Suppl. Fig. S9a). We quantified the expression level of 14,185 known NMD transcripts annotated by Ensembl release 101. After removing the non-expressed transcripts from the both flow cell runs, the remaining 6,423 NMD transcripts have shown significantly higher NMD transcript expression in the *SMG7*-KO/*SMG6*-KD (GFP-) cells than the WT (GFP+) cells (Suppl. Fig. S9b). We reason that the lowered NMD response is clearly visible by the enrichment of PTC-containing transcripts in the pool of *SMG7*-KO/*SMG6*-KD (GFP-) cells. Consequently, the cell barcode assignments meet our “biological” expectations.

We investigated a well-established NMD target *SRSF2* in detail (Sureau et al., 2001). The wildtype isoforms are present in both GFP+/-cells, while in the GFP-cells, the PTC-containing isoforms are more abundant in the GFP-cells (Suppl. Fig. S10a). The view on the *SRSF2* genome locus confirmed the different splicing junctions between two cell types (Suppl. Fig. S10b). The inclusion of exon 3 (middle) is clearly favored GFP-cells.

## DISCUSSION

The current ecosystem of single-cell RNA-seq platforms is rapidly expanding, but robust solutions for single-cell and single-molecule full-length RNA sequencing are virtually absent. In our manuscript, we combined Oxford Nanopore single-molecule sequencing of 10x Genomics cDNA libraries and developed a novel software tool to arrive at single-cell, single-molecule, full cDNA length resolution. In contrast to Lebrigand et al. (2020), our Bayesian method for cell barcode assignment performs superior in situation of low sequencing saturation. We could track in a well-controlled setting, i.e. by using GFP labeled cells and strong transcriptome pertubations, full-length transcript information at a single-cell level. We have identified differential RNA splicing linked to NMD pathway activity across our cell population. Our high-throughput full-length RNA sequencing solution is a necessary step forward towards studying the complex life of mRNA on single-cell level. This opens up unprecedented opportunities in low saturation settings such as multiplexed CRISPR-based screens.

## MATERIALS AND METHODS

### Single cell samples preparation and experiment

We performed an experiment using two different Flp-In-T-REx-293 cell lines: the wild type cell line with stably integrated FLAG-emGFP and a *SMG7* knockout (KO) cell line (generated and established in Boehm et al. (2020)). Wild type cells (GFP+) were transfected with siRNA against Luciferase and the *SMG7* KO cells (GFP-) were transfected with an siRNA against *SMG6*. Two days after siRNA transfection, we mixed both cell types at a 1:1 ratio with a target of 2,000 cells in total. A cDNA library was prepared according to the 10x Genomics Chromium Single Cell 3’ Reagent Kit User Guide (v3 Chemistry) from the pool of cells. The final libraries contain the P5 and P7 primers. The P5 read contains 21-nt adaptor sequence, 16-nt cellular barcode, 12-nt UMI, and polyA-tail, followed by cDNA sequences.

### Illumina reads processing and identification of cellular barcodes

We used 10X Genomics Cell Ranger 3.1 (https://github.com/10XGenomics/cellranger) to map the Illumina reads onto the reference genome. In our NMD dataset, the DNA sequences of luciferase were appended to the reference genome, and therefore the GFP+ cells can be called from Cell Ranger. Cell Ranger also corrects the sequencing errors in the barcode and unique molecular identifier (UMI) sequences. Cell Ranger estimates the number of cells using a Good-Turing frequency estimation model (https://support.10xgenomics.com), and characterized the identified barcodes into the cell-associated and background-associated barcodes. We used the cell-associated barcode sequences as the cellular barcode whitelist in the following analyses. Our CellRanger analysis estimated 2,052 sequenced cells (Suppl. Table S2).

### Nanopore reads processing, mapping, and gene assignment

We sequenced the two independently prepared Nanopore libraries from the same cDNA on two Nanopore R9.4 GridION flow cells (FC1 and FC2). The base-calling of Nanopore reads was done using Guppy v3.3.3, resulting 13,126,013 and 11,923,896 reads, respectively. We aligned the Nanopore reads onto the corresponding reference genome using minimap2 v2.17 (Li, 2018) in the spliced alignment mode (-ax splice). The two Nanopore runs yielded 11,158,994 and 10,164,820 mappable reads, respectively. We further assigned gene names to Nanopore reads using the “TagReadWithGeneExon” program from the Drop-seq tools (Macosko et al., 2015). We assembled all the Nanopore reads and extended transcriptome annotations using StringTie v2.1.1 (Pertea et al., 2015). The FPKM level of the assembled transcripts were quantified using Ballgown v2.14.1 (Frazee et al., 2015).

### Identification of the adapter, barcode, UMI, and polyA-tail sequences from Nanopore reads

We removed the cDNA sequences from Nanopore reads, and extracted up to 100bp from both ends. We developed a modified version of FLEXBAR (Dodt et al., 2012; Roehr et al., 2017) to align P1 primer adapter sequence with the following parameters (“-ao 10 -ae 0.3-ag -2 -hr T -hi 10 -he 0.3 -be 0.2 -bg -2 -bo 5 -ul 26 -kb 3 -fl 100”). Then we aligned the Nanopore reads that have valid adapters to the cellular barcodes which have been previously identified by Cell Ranger. We scanned the poly-A sequences using the homopolymer-trimming function of FLEXBAR downstream of the cell barcode. Once the poly-A sequences were found, the UMI sequences between the poly-A and barcode were searched using MUMmer 4.0 (Marçais et al., 2018) (with parameters “-maxmatch -b -c -l 7 -F”) and in-house scripts against the Illumina UMIs of the same cell and the same gene or genomic regions (± 500bp from each end of the reads). In the end, ScNapBar output the alignment score of the adapter, the number of mismatches and indel from the barcode alignment, the length of poly-A and UMI sequences, as well as the length of the gap between the barcode and adapter. We use these features to estimate the likelihood of the barcode assignment in the following steps (Fig. 1).

### Simulation and engineering of discriminative features from the barcode and adapter alignments

We characterized the correct and false barcode assignment by simulating Nanopore reads. We created some artificial template sequences which contain only the P1 primer, cellular barcode, and UMI sequences at the same frequencies as the Illumina library, followed by 20bp oligo-dT and 32bp cDNA sequences. In the next step, we first used NanoSim (Yang et al., 2017) to estimate the error profile of our Nanopore library, then we generated one million Nanopore reads from the artificial template using the NanoSim simulator with the previously estimated error profile. We aligned the simulated Nanopore reads to the adapter and barcode sequences using ScNapBar. We compared the sequences in the simulated Nanopore reads and the sequences from the artificial template, and labeled the assigned barcode as correct or false accordingly. By comparing sequence and alignment features of correct and false assignments, we found that the two categories (false, true) could be discriminated by these features (Suppl. Fig. S8c). We then assessed the importance of each feature towards the correctness of the assignment (Suppl. Fig. S8a). As these features are uncorrelated (Suppl. Fig. S8b), we train a Naïve Bayes model from these features to predict the likelihood of the correctness of a barcode assignment.

### Calculate cell barcode posterior probability using prior probabilities from the Illumina data set

We denote *b*_1_,*b*_2_,…,*b_n_* as barcodes that match to read *r* and define *P*(*b*_1_|*r*) as the probability that barcode *b*_1_ was sequenced given *r* is observed. Following Bayes’ theorem, *P*(*b*_1_|*r*) could be computed as in Eq. 3a, and further computed as in Eq. 3b according to the total probability theorem.

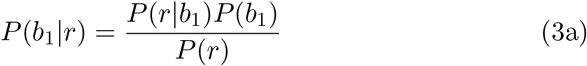

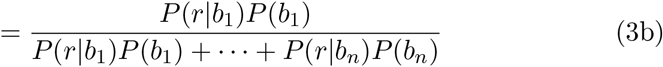

where *P*(*r|b*_1_) and *P*(*r|b_n_*) are computed by the Naïve Bayes predictor, and priors *P*(*b*_1_) and *P*(*b_n_*) can be estimated from the observed barcode counts in Illumina sequencing. For practical reasons, as the probabilities for the unaligned barcodes that contain a lot of mismatches are pretty low, we add a pseudocount of 1 to the denominator to represent them. Because we have sequenced the same library twice using the Nanopore and Illumina sequencer, we assume prior probabilities *P*(*b*) are the same for the Nanopore and the Illumina platform (Suppl. Fig. S2a).

### Quality assessment and clustering of the single-cell libraries

A meta gene body coverage analysis confirmed the near full-length character of the Nanopore approach (Suppl. Fig. S6a). After assigning gene names and cell barcodes to the Nanopore reads, we processed the gene-barcode expression matrix using Seurat v3.1.1 (Butler et al., 2018) by keeping the genes expressed in minimal three cells, and cells with more than 200 genes expressed. We then scaled the expression matrix by a factor of 10,000 and log-normalized, and performed the t-SNE analysis.

## DATA DEPOSITION

All sequencing data were deposited in NBCI’s SRA database (accession number). ScNapBar workflow (code and tutorial) is available at https://github.com/dieterich-lab/single-cell-nanopore.

## AUTHOR’S CONTRIBUTIONS

QW implemented the ScNapBar workflow, performed data analyses and wrote the manuscript. SB modified the FLEXBAR implementation. VB performed cell culture experiments and helped to draft the manuscript. NG helped to draft the manuscript and acquired funding. JA performed all sequencing experiments and helped to draft the manuscript. CD supervised the project, performed data analyses, acquired funding and wrote the manuscript.

## ACKNOWLEDGMENTS

The authors would like to thank Etienne Boileau, Thiago Britto Borges, Tobias Jakobi for proof-reading and comments. The authors are grateful to Marek Franitza for running the experiments on the 10x platform and to Christian Becker for running ONT sequencing. Supported by Informatics for Life funded by the Klaus Tschira Foundation.

